# cryoSPHERE: Single-particle heterogeneous reconstruction from cryo EM

**DOI:** 10.1101/2024.06.19.599686

**Authors:** Gabriel Ducrocq, Lukas Grunewald, Sebastian Westenhoff, Fredrik Lindsten

## Abstract

The three-dimensional structure of a protein plays a key role in determining its function. Methods like AlphaFold have revolutionized protein structure prediction based only on the amino-acid sequence. However, proteins often appear in multiple different conformations, and it is highly relevant to resolve the full conformational distribution. Single-particle cryo-electron microscopy (cryo EM) is a powerful tool for capturing a large number of images of a given protein, frequently in different conformations (referred to as *particles*). The images are, however, very noisy projections of the protein, and traditional methods for cryo EM reconstruction are limited to recovering a single, or a few, conformations. In this paper, we introduce cryoSPHERE, a deep learning method that takes as input a nominal protein structure, e.g. from AlphaFold, learns how to divide it into segments, and how to move these as approximately rigid bodies to fit the different conformations present in the cryo EM dataset. This formulation is shown to provide enough constraints to recover meaningful reconstructions of single protein structures. This is illustrated in three examples where we show consistent improvements over the current state-of-the-art for heterogeneous reconstruction.

## 1. Introduction

Single-particle cryo-electron microscopy (cryo EM) is a powerful technique for determining the three-dimensional structure of biological macromolecules, including proteins. In a cryo EM experiment, millions of copies of the same protein are first frozen in a thin layer of vitreous ice and then imaged using an electron microscope. This yields a micrograph: a noisy image containing 2D projections of individual proteins. The protein projections are then located on this micrograph and cut out so that an experiment typically yields 10^4^ to 10^7^ images of size N_pix_ × N_pix_ of individual proteins, referred to as *particles*. Our goal is to reconstruct the possible structures of the proteins given these images.

Frequently, proteins are conformationally heterogeneous and each copy represents a different structure. Conventionally, this information has been discarded, and all of the sampled structures were assumed to be in only one or a few conformations (*homogeneous* reconstruction). Here, we would like to recover all of the structures in a *heterogeneous* reconstruction.

Structure reconstruction from cryo EM presents a number of challenges. Firstly, each image shows a particle in a different, unknown orientation. Secondly, because of the way the electrons interact with the protein, the spectrum of the images is flipped and reduced. Mathematically, this corresponds to a convolution of each individual image with the Point Spread Function (PSF). Thirdly, the images typically have a very low signal-to-noise ratio (SNR), usually around 0.1, see e.g. Baxter et al. (2009).

For all these reasons, it is very challenging to perform *de novo* cryo EM reconstruction. Standard methods, produce electron densities averaged over many, if not all conformations (Scheres, 2012; Punjani et al., 2017), performing discrete heterogeneous reconstruction. More recent methods attempt to extract continuous conformational heterogeneity, e.g. by imposing constraints on the problem through an underlying structure deformed to fit the different conformations present in the dataset, see e.g Rosenbaum et al. (2021); Zhong et al. (2021b); Li et al. (2023). AlphaFold (Jumper et al., 2021) and RosettaFold (Baek et al., 2021) can provide such a structure based on the primary sequence of the protein only. In spite of this strong prior, it is still difficult to recover meaningful conformations. The amount of noise and the fact that we observe only 2D projections creates local minima that are difficult to escape (Zhong et al., 2021b; Rosenbaum et al., 2021), leading to unrealistic conformations.

To remedy this, we root our method in the observation that different conformations can often be explained by large scale movements of domains of the protein (Mardt et al., 2022). Specifically, we develop a variational auto-encoder (VAE) (Kingma & Welling, 2014) that, from a nominal structure and a set of cryo EM images:

- Learns how to divide the amino-acid chain into segments, given a user defined maximum number of segments; see Figure 2. The nominal structure can for instance be obtained by AlphaFold (Jumper et al., 2021).
- For each image, learns approximately rigid transformations of the identified segments of the nominal structure, which effectively allows us to recover different conformations on an image-by-image (single particle) basis.

These two steps happen concurrently, and the model is end-to-end differentiable. The model is illustrated in Figure 1.

**Figure 1.**
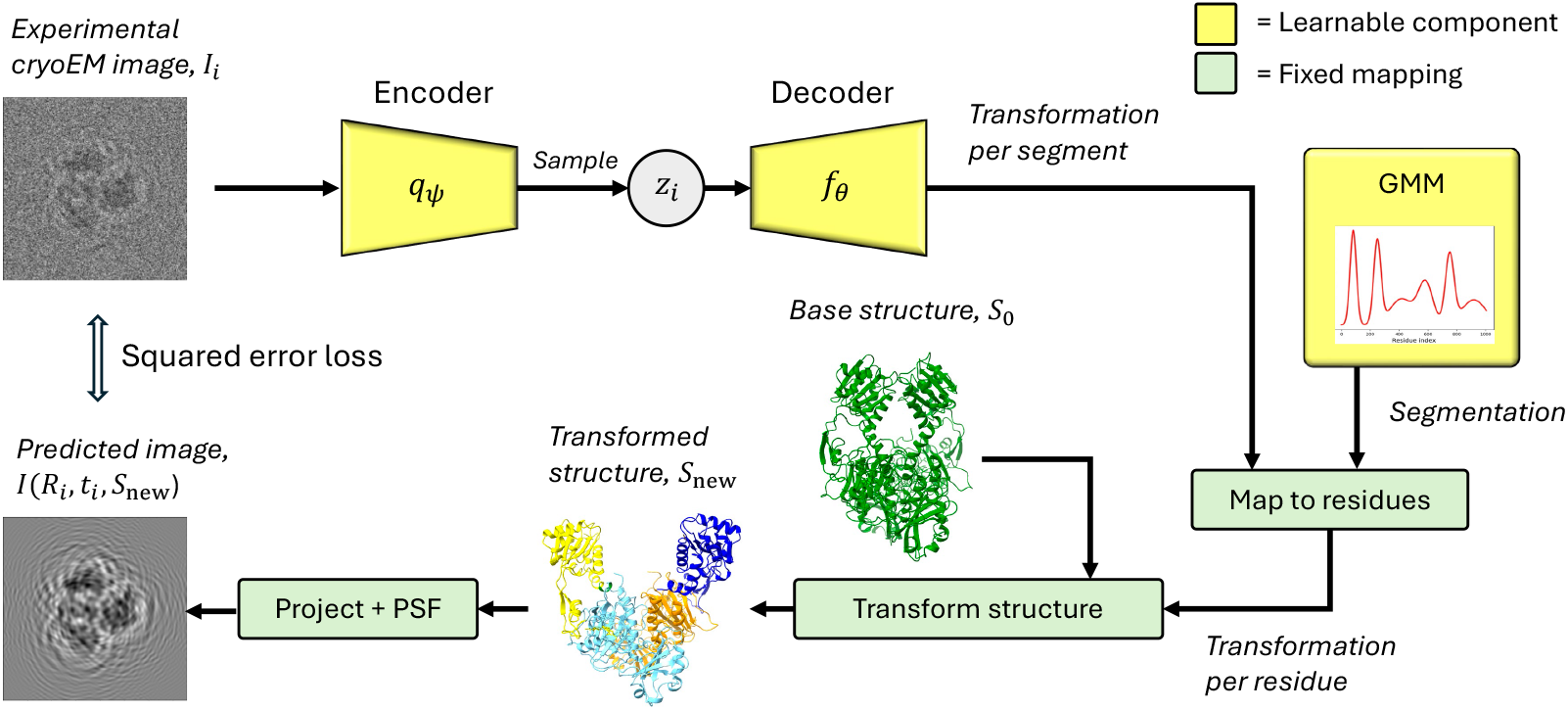
Flow chart of our network. The learnable parts of the model are the encoder, the decoder and the Gaussian mixture. Note that even though the transformations predicted by the decoder are on a per image basis, that is not the case of the Gaussian mixture, which is shared across all particles.

**Figure 2.**
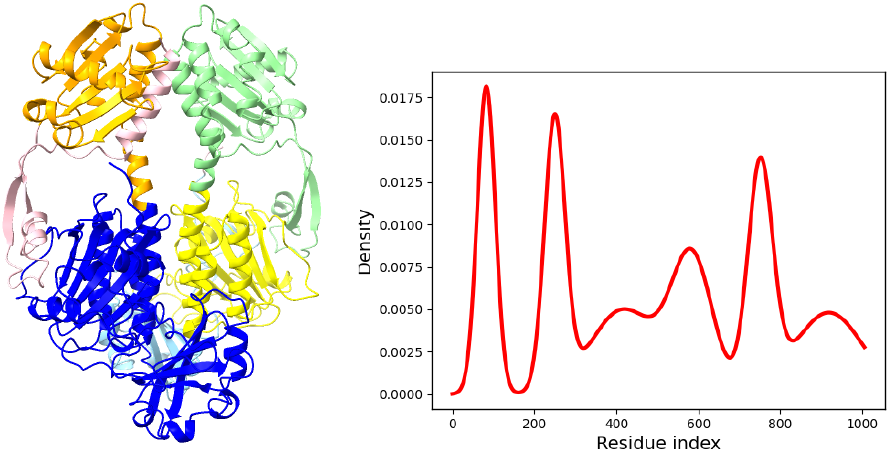
Example of segments recovered with a Gaussian mixture of 6 components.

Note that what we call a segment is conceptually different from a domain in the structural biology sense. The domains of a protein play a pivotal role in diverse functions, engaging in interactions with other proteins, DNA/RNA, or ligand, while also serving as catalytic sites that contribute significantly to the overall functionality of the protein, see e.g. Schulz & Schirmer (1979); Nelson et al. (2017). By comparison, the segments we learn do not necessarily have a biological function. However, while not strictly necessary for the function of the method, experiments in Section 4 show that our VAE often recovered the actual domains corresponding to different conformations.

The implementation of the model is available on github ^1^.

## 2. Background

### 2.1. Notations and problem formulation

In what follows, *S* = {*r*_*i*_}_*i*_ = 1,…,*R*_res_ denotes a protein made of a number *R*_res_ ∈ ℕ^⋆^ of residues *r*_*i*_. Each residue is itself a set of atoms : 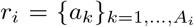 where *A*_*i*_ ∈ ℕ^⋆^ is the number of atoms present in residue *i*. We can equivalently define the structure 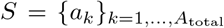 in terms of its atoms, where *A*_total_ ∈ ℕ^⋆^ is the total number of atoms in the structure. We denote by *a*_*k*_ = (*x*_*k*_, *y*_*k*_, *z*_*k*_, *η*_*k*_)^*T*^ ∈ ℝ_3_ × {*C*_*α*_, *C*_*β*_, *N*} the coordinates and atom type of *a*_*k*_ for *k* ∈ {1, …, *A*_total_}. This means that in this paper, we limit ourselves to the backbone of the protein: we consider only the *C*_*α*_, *C*_*β*_ and *N* atoms.

The electron density map of a structure S, also called a volume, is a function *V*_*S*_ : ℝ^3^ → ℝ, where *V*_*S*_(*x*) represents the probability density function of an electron of *S* being present, evaluated at x ∈ ℝ^3^. That is, if B ⊆ ℝ^3^, the probability that the structure *S* has an electron in *B* is given by ∫_*B*_*V*_*S*_*(x)dx*.

Assume we have a set of 2D images 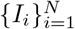 of size *N*_pix_ × *N*_pix_, representing 2D projections of different copies of the same protein in different conformations. Traditionally, the goal of cryo EM heterogeneous reconstruction has been to recover, for each image *i*, the electron density map *V*_*i*_ corresponding to the underlying conformation present in image *i*. See Subsection 2.2 for a review these methods. However, following recent works, e.g., Rosenbaum et al. (2021); Zhong et al. (2021b), we aim at recovering, for each image *i*, the underlying structure *S*_*i*_ explaining the image. That is, we try to recover the precise position in ℝ^3^ of each residue.

Since the different conformations of large proteins can often be explained by large scale movements of its domains (Mardt et al., 2022), our method learns to decompose the amino-acid chain of the protein into segments and, for each image *I*_*i*_, to rigidly move the learnt segments of a base structure *S*_*0*_ to match the conformation present in that image, as explained in Section 1.

AlphaFold (Jumper et al., 2021) and RosettaFold (Baek et al., 2021) can provide such a structure *S*_*0*_, based on the amino-acid sequence of the protein at hand. In Section 4, we further fit the AlphaFold predicted structure into a volume recovered by a custom backprojection algorithm provided by Zhong et al. (2020).

### 2.2. Related work

Two of the most popular methods for cryo EM reconstruction, which are *not* based on deep learning, are RELION (Scheres, 2012) and cryoSPARC (Punjani et al., 2017). Both methods perform volume reconstruction, hypothesize that *k* conformations are present in the dataset and perform maximum a posteriori estimation over the *k* density maps, thus performing discrete heterogeneous reconstruction. Both of these algorithms operate in Fourier space using an expectation-maximization algorithm (Dempster et al., 1977) and are non-amortized: the poses are learnt for each image, and adding new images to the dataset would require a new run of the algorithm. Other approaches perform continuous heterogeneous reconstruction. For example, 3DVA (Punjani & Fleet, 2021b) uses a probabilistic principal component analysis model to learn a latent space while E2GMM (Chen & Ludtke, 2021) parameterizes the volume as a mixture of *N* Gaussian p.d.f and learns their mean conditionally on an input image.

Another class of methods involve deep learning and typically performs continuous heterogeneous reconstruction using a VAE architecture. Some of them attempt to reconstruct a density map, while others target the atomic structures – or a backbone – directly. For example, cryoDRGN (Zhong et al., 2020; 2021a) and CryoAI (Levy et al., 2022) use a VAE acting on Fourier space to learn a latent space and a mapping that associates a 3D density map with each latent variable. They perform non-amortized and amortized inference over the poses, respectively. Other methods are defined in the image space, e.g. 3DFlex (Punjani & Fleet, 2021a) and cryoPoseNet (Nashed et al., 2021). They both perform non-amortized inference over the poses. These methods either learn, for a given image *I*_*i*_, {*V*_*i*_(*x*_*k*_)} the values at a set of 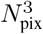 fixed 3D coordinates {*x*_*k*_}, representing the volume on a grid (*explicit* parameterization), or an actual function 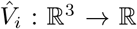 in the form of a neural network that can be queried at chosen coordinates (*implicit* parameterization). The implicit parameterization requires smaller networks but needs an expensive 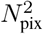 and 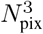 evaluation for image and volume generation, respectively.

Other deep learning methods attempt to directly reconstruct structures instead of volumes and share a common process: starting from a plausible base structure, obtained with e.g. AlphaFold (Jumper et al., 2021), for each image, they move each residue of the base structure to fit the conformation present in that specific image. These methods differ on how they parameterize the structure and in the prior they impose on the deformed structure or the motion of the residues. For example AtomVAE (Rosenbaum et al., 2021) considers only residues and penalizes the distances between two subsequent residues that deviate too much from an expected value. CryoFold (Zhong et al., 2021b) considers the residue centers and their side-chain and also imposes a loss on the distances between subsequent residues and the distances between the residue centers and their side-chain. Unfortunately, due to the high level of noise and the fact that we observe only projections of the structures, these ”per-residue transformation” methods tend to be stuck in local minima, yielding unrealistic conformations unless the base structure is taken from the distribution of conformations present in the images (Zhong et al., 2021b), limiting their applicability to real datasets. Even though AtomVAE (Rosenbaum et al., 2021) could roughly approximate the distribution of states of the protein, it was not able to recover the conformation given a specific image. In addition, to our knowledge, the code for these two methods is not available and non-trivial to reimplement. This prevents a comparison of our results with AtomVAE and CryoFold.To reduce the bias that the base structure brings, (Schwab et al., 2023) recently proposed to fit pseudo-atoms in a consensus map with a neural network directly. Similar to our work, several other methods constrain the atomic model to rigid body motions. For example (Chen et al., 2023), starting from a base structure *S*_big_ fitted in the consensus map (called the big GMM), learn a representation structure *S*_small_ of this consensus map using a smaller number using the method described in (Schwab et al., 2023). They subsequently learn a translation on each residue *S*_small_, that contribute to translate the residues of *S*_big_ weighted by how close they are from the residues of *S*_small_. This can be thought of as our segmentation GMM, except that it takes places in the 3D space and only the centers of the Gausian modes can be learnt, while ours is fitted on the residue indexes and all of the GMM parameters are learnable. In addition, they do not use this segmentation to perform rigid body motions. Instead, in a subsequent step, the user is in charge of identifying the rigidly moving segments of the protein by applying one or several masks to the protein. This is in contrast to our method where the segmentation used to perform rigid body motion is learnt by the network trying to move the domains to fit the images. Finally, this method involves a sequence of optimization problem, with potentially different architecture, while ours is end-to-end differentiable. Using the method in (Schwab et al., 2023), Chen et al. (2024) develop a focused refinement on patches of the GMM representation of the protein. These patches are learnt using k-means on the location of residues and do not depend on the different conformations of the data set. This in contrast to cryoSPHERE which learns the segments of the protein are tightly linked to the change of conformation. Concurrently to our work, Li et al. (2023) developed CryoSTAR which acts on each residue and enforces local rigidity of the motion of the protein by imposing a similarity loss between the base structure and the deformed structure. The interested reader can see Donnat et al. (2022) for an in-depth review of deep learning methods for cryo EM reconstruction.

The reconstruction methods relying on an atomic model, such as e.g cryoSTAR, DynaMight or cryoSPHERE offer the possibility to the user to provide prior information via this atomic model. They also offer the possibility of deforming the protein according to chemical force fields. This is not the case of the methods performing volume reconstruction without such an atomic model.

## 3. Method – cryoSPHERE

In this section, we present our method for single-particle heterogeneous reconstruction, denoted cryoSPHERE. The method focuses on structure instead of volume reconstruction. It differs from the previous works along this line in the way the movements of the residues are constrained: instead of deforming the base structure on a residue level and then imposing a loss on the reconstructed structure, it moves parts of the protein approximately rigidly by construction.

We use a typical VAE architecture, see Figure 1 for a depiction of how the information flows in our network. We map each image to a latent variable by a stochastic encoder. The latent variable is then passed to a decoder which outputs a transformation (rotation and translation) per segment. Based on these transformations and segment decomposition, the underlying structure *S*_0_ is deformed and then turned into a volume. Finally, the volume is projected into a 2D image, according to its pose that can be estimated using existing software, e.g. (Scheres, 2012; Punjani et al., 2017). Finally, this image is convolved with the PSF to give the reconstructed image that is compared to the input image. This completes the forward pass. In the backward pass, the parameters of the encoder, decoder and Gaussian mixture are updated with a gradient descent based on the evidence lower bound (see Section 3.2). In the following, we describe details of our network: the encoder and decoder, the segment decomposition, how to turn a structure into a 2D projection of its electron density, and an efficient way to compare the true and predicted images.

### 3.1. Image formation model

To compute the 2D projection of the protein structure *S*, we first estimate its 3D electron density map *V* :

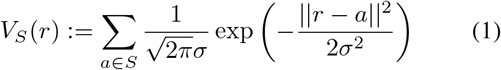

where r ∈ ℝ^3^ and *σ* is a positive real number. In this definition the protein’s electron density is approximated as the sum of a Gaussian p.d.f centered on each atom. We use a fixed value of σ = 1, following Rosenbaum et al. (2021).

From these density maps, we then compute an image projection 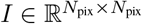 as:

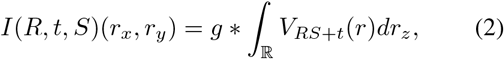

where (*r*_*x*_, *r*_*y*_) ∈ ℝ^2^ are the coordinates of a pixel, *r*_*z*_ ∈ ℝ is the coordinate along the *z* axis, *R* ∈ *SO*(3) is a rotation matrix and *t* ∈ ℝ^3^ is a translation vector. The abuse of notation *RS* + *t* means that every atom of *S* is rotated according to *R* and then translated according to *t*. The image is finally convolved with the point spread function (PSF) *g*, which in Fourier space is the contrast transfer function (CTF), see Vulović et al. (2013).

In practice, we avoid approximating the integral in Equation (2). The density map, being a sum of Gaussian p.d.f, allows for an exact calculation by integrating over the last marginal. This significantly reduces the computing time.

### 3.2. Maximum likelihood with variational inference

To learn a distribution of the different conformations, we hypothesize that the conformation seen in image *I*_*i*_ depends on a latent variable *z*_*i*_ ∈ ℝ^*L*^, with prior *p*(*z*_*i*_). Let *f*_*θ*_(*S*_*0*_, *z*) be a function which, for a given base structure *S*_*0*_ and latent variable *z*, outputs a new transformed structure *S*. This function depends on a set of learnable parameters *θ*. Then, the conditional likelihood of an image 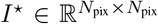 with a pose given by a rotation matrix *R* and a translation vector t is modeled as 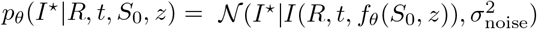, where 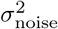 is the variance of the observation noise. The marginal likelihood is thus given by

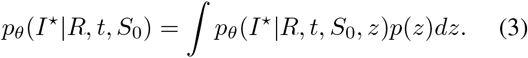

In practice, the pose (*R, t*) of a given image is unknown. However, following similar works (Zhong et al., 2021b; Li et al., 2023), we suppose that we can estimate R and t to sufficient accuracy using off-the-shelf methods (Scheres, 2012; Punjani et al., 2017).

Directly maximizing the likelihood (3) is infeasible because one needs to marginalize over the latent variable. For this reason, we adopt the VAE framework, conducting variational inference on *p*_*θ*_(*z*|*I*^*⋆*^) ∝ *p*_*θ*_(*I*^*⋆*^|*z*)*p*(*z*), and simultaneously performing maximum likelihood estimation on the parameters *θ*.

Let *q*_*ψ*_(*z* |*I*^*⋆*^) denote an approximate posterior distribution over the latent variables. We can then maximize the evidence lower-bound (ELBO):

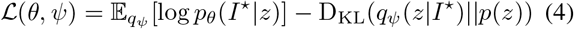

which lower bounds the log-likelihood log *p*_*θ*_(*I*^*⋆*^). Here D_KL_ denotes the Kullback-Leibler (KL) divergence. In this framework *f*_*θ*_ is called the decoder and *q*_*ψ*_(*z*|*I*^⋆^) the encoder.

### 3.3. Segment decomposition

We fix a maximum number of segments *N*_segm_ ∈ {1, …, *R*_*res*_} and we represent the decomposition of the protein by a stochastic matrix 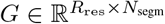. The rows of *G* represent ”how much of each residue belongs to each segment”, and our objective is to ensure that each residue mostly belongs to one segment, that is:

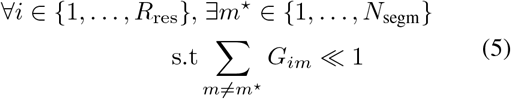

We also aim for the segments to respect the sequential structure of the amino acid chain, and the model to be end-to-end differentiable. Without end-to-end differentiability, we could not apply the reparameterization trick and we would have to resort on Monte-Carlo estimation of the gradient of the segments, which has a higher variance, see e.g (Mohamed et al., 2019).

To meet these criteria, we fit a Gaussian mixture model (GMM) with *N*_*segm*_ components on the real line supporting the residue indices, that is, on ℝ. Each component *m* has a mean *µ*_*m*_, standard deviation *σ*_*m*_ and a softmax weight *α*_*m*_. The {*α*_*m*_} are passed into a softmax to obtain the weights {*π*_*m*_} of the GMM, ensuring they are positive and summing to one. We define the probability that a residue *i* belongs to segment m as the probability that the index *i* belongs to mode m, annealed by a temperature τ ∈ ℝ^+⋆^:

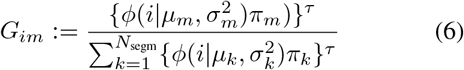

where *ϕ*(*x* | *µ, σ*^*2*^) is the unidimensional Gaussian probability density function with mean *µ* and variance *σ*^2^ and τ is a fixed hyperparameter. If τ is sufficiently large, we can expect condition (5) to be verified. See Figure 2 for an example of a segment decomposition using a Gaussian mixture.

In this “soft” decomposition of the protein, each residue can belong to more than one segment, allowing for smooth deformations. In addition, the differentiable architecture is amenable to gradient descent methods, and a well chosen τ can approximate a “hard” decomposition of the protein. We set τ = 20 in the experiment section. In our experience, this segmentation procedure is very robust to different initialization and converge in only a few epochs.

### 3.4. Decoder architecture

The decoder describes the distribution of the images given the latent variables. The latent variables include:

1. One latent variable *z*_*i*_ ∈ *ℝ*^*L*^ per image, parameterizing the conformation.
2. The global parameters {*µ*_*m*_, *σ*_*m*_, *α*_*m*_} _*m=1*,…,*Nsegm*_ of the GMM describing the segment decomposition.

Given these latent variables and a base structure *S*_*0*_, we parameterize the decoder *f*_*θ*_ in three steps.

First, a neural network with parameters *θ* maps *z*_*i*_ ∈ *ℝ*^*L*^ to one 3D translation per segment 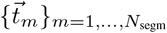 and one rotation per segment. The rotation is represented by a unit quaternion 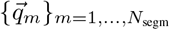 (Vicci, 2001), which can be equivalently described as a set of axes of rotation 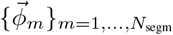 and corresponding angles of rotations 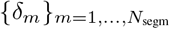.

Second, given the parameters of the GMM, we can compute the matrix *G*. Finally, for each residue *i* of *S*_*0*_, we update the coordinates of all its atoms {*a*_*ik*_}_*k=1*,…,*Ai*_ :

1. First, *a*_*ik*_ is successively rotated around the axis 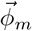 with an angle *G*_*im*_*δ*_*m*_ for *m* ∈ {1, …, *N*_*segm*_} to obtain updated coordinates *a*^*′*^_*ik*_.
2. Second, it is translated according to:

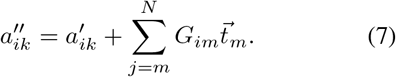

This way, the rotation angles and translations for a residue incorporate contributions from all segments, proportionally on how much they belong to the segments. If condition (5) is met, a roughly rigid motion for each segment can be expected.

### 3.5. Encoder and priors

We follow the classical variational-autoencoder framework. The distribution *q*_*ψ*_(*y* | *I*^*⋆*^) is given by a normal distribution 𝒩(*µ*(*I*^*⋆*^), diag(*σ*^*2*^(*I*^*⋆*^))) where *µ* ∈ ℝ^L^ and 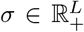 are generated by a neural network with parameters *ψ*, taking an image *I*^*⋆*^ as input. Additionaly, the approximate posterior distribution on the parameters of the GMM is chosen to be Gaussian and independent of the input image:

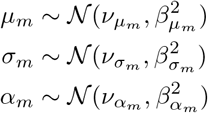

where 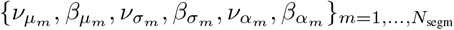 are parameters that are directly optimized and in practice we feed *σ*_*m*_ to an ELU+1 layer in order to avoid a negative or null standard deviation.

Finally, we assign standard Gaussian priors to both the local latent variable *z*_*i*_ *z*_*i*_ ∼ 𝒩 (0, *I*_*L*_), and the global GMM parameters {*µ*_*m*_, *σ*_*m*_, *α*_*m*_, }_*m*=1,…,*Nsegm*_.

Note that applying the reparameterization trick (Kingma & Welling, 2014) is straightforward for a Gaussian distribution. Calculating the KL-divergence between two Gaussian distributions as in Equation (4), is also straightforward.

## 4. Experiments

In this section, we test cryoSPHERE on different datasets and compare the results to cryoDRGN (Zhong et al., 2020). CryoDRGN is a state-of-the-art method for continuous heterogeneous reconstruction, in which the refinement occurs at the level of electron densities. To synthesize data for all experiments, we applied CTF corruption to the images and add Gaussian noise to achieve SNR ≈ 0.1. See Appendix A for a detailed description of the CTF. Furthermore, we make the assumption that we know the exact poses. However, since we use a structure S_0_, which is different from the structures used to generate the datasets, these poses can only be an approximation. This will not be the case of cryo-DRGN, for which these poses will indeed be exact, as this method does not use a base structure. Consequently, in this context, the comparison may introduce a bias in favour of cryoDRGN.

### 4.1. Toy dataset: a continuous motion

For an initial assessment of cryoSPHERE, we examine a simulated dataset based on a bacterial phytochrome with UniProt (The UniProt Consortium, 2021) entry Q9RZA4. This protein is a dimer, with 755 residues per chain. To simulate a continuous heterogeneous set of images, we predicted the structure with AlphaFold multimer (Evans et al., 2021) and divided the protein into two domains. The first one consists of the full chain A and 598 residues of chain B, 1353 residues in total. The second domain comprises the last 157 residues of the chain B, see Figure 3. We then sample 10^4^ rotation angles from a two-components Gaussian mixture. Each conformation is generated by rotating the second domain according to the axis (0, 1, 0) and one of the sampled rotation angles. This yields 10^4^ different conformations. Finally, we uniformly sample 15 different poses for each structure, yielding a total of 150k images. Appendix A.1 provides more details, and Figure 3 shows examples of structures.

**Figure 3.**
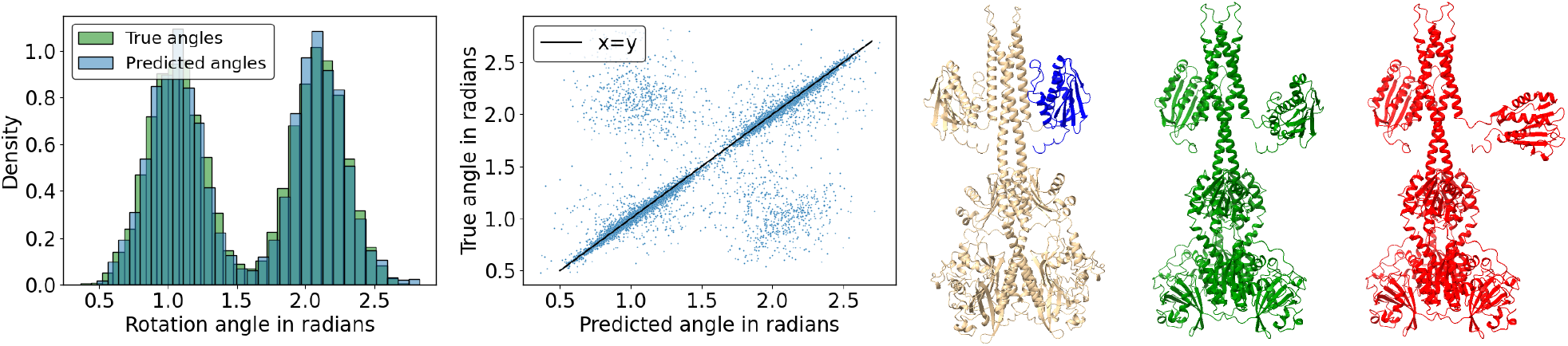
Toy dataset. Leftmost: Histograms of the predicted and true angles of rotation in radians. The true angles are in green. The recovered distances are in blue. Left: Predicted against true angles in å ngström. The black line represent *x* = *y*. Middle: Base structure with the two domains, also used to generate the images. The domain in blue is rotated according to the axis (0, 1, 0) with angles of rotations sampled from the green distribution on the leftmost figure. Note that the segments predicted by cryoSPHERE exactly match the two domains. The fourth segment is in blue and corresponds to the last 157 residues of chain B, matching exactly the ground truth domain. Right: First mode structure. Rightmost: Second mode structure.

Testing the segment decomposition, we then run cryoSPHERE by requesting division into *N*_*segm*_ = 4. The program learnt a first and third segment with 0 residues, a second segment with 1353 residues and a fourth segment with 157 residues (Figure 3). Thus, cryoSPHERE learnt segments according to the ground truth.

Moreover, Figure 3 shows that most of the predicted angles of rotation for the fourth segment are in excellent agreement with the ground truth structural changes. However, approximately 12 % of the dataset is predicted as belonging to the wrong mode. Figure 8 in the appendix shows that the norms of the predicted translations for both segments are negligible, also indicating that cryoSPHERE has recovered the ground truth. The same figure shows that we recover the right axis of rotation. In Appendix A.1, we plot the predicted angles against the mean of the latent variable distributions output by the encoder. We observe that cryoSPHERE has recovered the circular motion: the greater the latent mean, the lower the predicted angle, with a quick transition from one mode to the other. This indicates that cryoSPHERE is able to recover the structure of single-particles from cryo EM data.

### 4.2. A discrete conformation dataset: the SARS-CoV-2 spike protein

We now test cryoSPHERE on a dataset comprising two discrete conformations of the SARS-CoV-2 spike protein: a close conformation with PDB entry 6ZP5 and an open conformation with PDB entry 6ZP7, see Melero et al. (2020). We generate 150k images of size *N*_pix_ = 300 with uniformly sampled poses and neglected translations, with seventy percent corresponding to the close conformation and thirty percent to the open one.

Prior to running cryoSPHERE, we obtain a base structure by first running the multimer version of AlphaFold (Evans et al., 2021) on the amino acid sequence of the spike protein, selecting the best (according to AlphaFold) structures. Second, we utilize a custom backprojection algorithm provided by (Zhong et al., 2020) to get a homogeneous reconstruction. This yields a volume with low resolution in the mobile parts but decent resolution in the less mobile ones. To reduce a potential bias introduced by using an AlphaFold structure, we subsequently fit this structure into the volume, initially performing rigid-body fitting in ChimeraX (Meng et al., 2023) and then using ISOLDE (Croll, 2018). Subsequently, we run cryoSPHERE with this fitted structure as the base structure for 48 hours on a single GPU, totalling 174 epochs.

We define the two conformations in terms of the angle between two domains, see Figure 4. The same figure demonstrates that we recover the angles within roughly 1 degree. After clustering the structures predicted by cryoSPHERE based on their angle defined in Figure 4, only 4 of them get predicted to be in the wrong conformation. Additionally, by turning the structures predicted by cryoSPHERE into volumes using our operator (1), we observe excellent agreement with the ground truth volumes, as shown by the Fourier shell correlation (FSC) curves (Klaholz, 2015). This illustrates that cryoSPHERE can effectively recover discrete conformational disorder.

**Figure 4.**
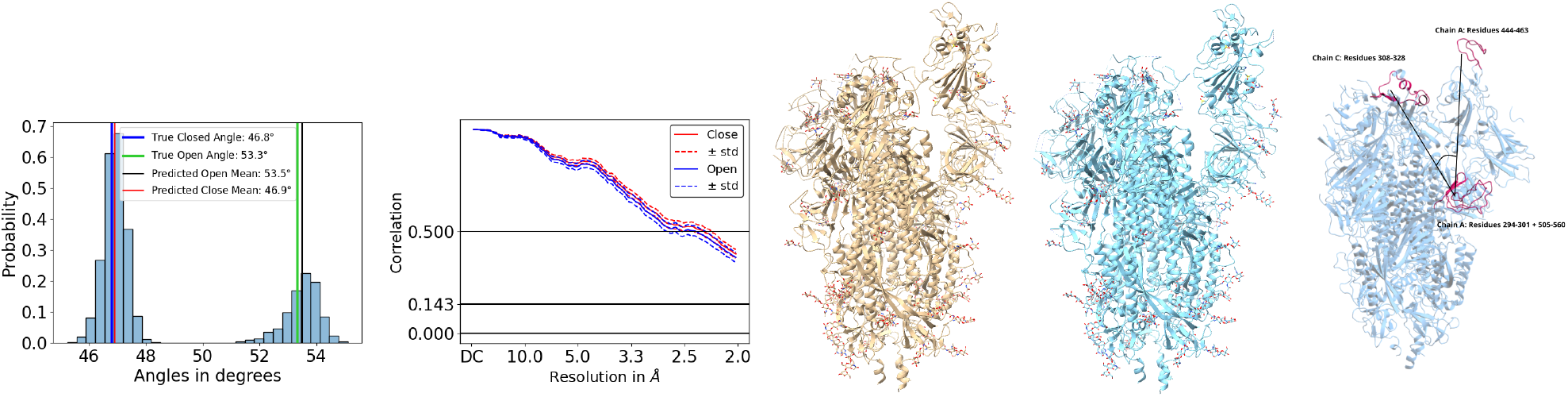
Spike dataset. Left: Predicted angles. Middle left: FSC curves for the open and close conformations. Middle: Close conformation. Middle right: Open conformation. Right: Definition of the angles.

### 4.3. Molecular dynamics dataset: bacterial phytochrome

We now test cryoSPHERE with a more difficult dataset. This time, we simulate a continuous motion of a bacterial phytochrome, with PDB entry 4Q0J (Burgie et al., 2014). This dimer has 503 residues per chain. We define two upper domains corresponding to the residues 321-502 from both chain A and B. Then, we run a Molecular Dynamics (MD) simulation to simulate a dissociation of these two domains, as depicted in Figure 9 in Appendix A.2. We take 10^4^ structures along the trajectory and sample 15 poses uniformly for each structure. Refer to Appendix A.2 for more details.

Analogously to Section 4.2, we obtain an AlphaFold structure that we fit into a volume reconstructed by backprojection. Further details can be found in Appendix A.2. Finally, we run our method with the fitted AlphaFold structure as our base structure and *N*_*segm*_ = 20.

Since cryoSPHERE outputs structures, we can compute the predicted distances between the above-mentioned domains. We plot the true and recovered histograms of the distances between the two domains in Figure 5. Additionally, we plot the predicted distances against the true distances in the same figure. These two plots show that the predicted structures are in excellent agreement with the ground truth structures. This is even true for the most dissociated structures, even though the majority of the structures are in a closed conformation. Figure 6 shows the recovered segments as well as a selection of predicted structures and the corresponding ground truths. Our model has learnt to separate the two mobile top domains from the fix bottom one.

**Figure 5.**
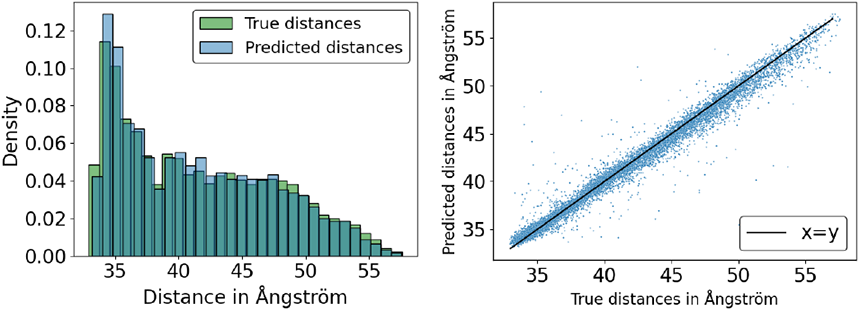
MD dataset. Left: Histograms of the distances of the two upper domains. The true distances are in green. The recovered distances are in blue. Right: Predicted against true distances in å ngström. The black line represent *x* = *y*.

**Figure 6.**
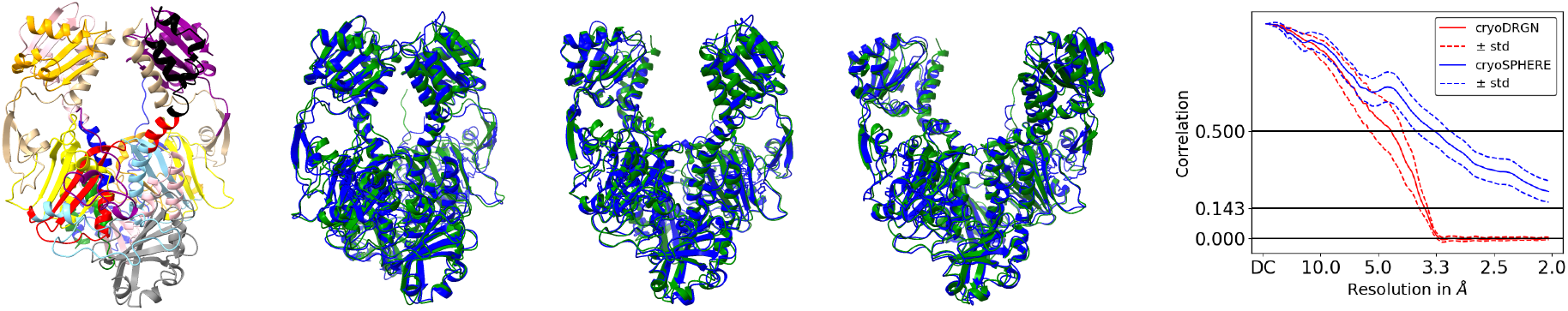
MD dataset. Left: Recovered segments on the fitted AlphaFold structure. The colors denotes different contiguous domains: if two segments share the same color, it does not mean their residues belong to each others. Middle left to middle right: cryoSPHERE predicted structures in green, corresponding ground truth in blue. Right: Fourier shell correlation curves. The mean across structures for cryoSPHERE is in plain blue. Dotted blue represents the mean *±* one standard deviation. The red, plain line represents the mean for cryoDRGN across volumes, with the dotted red line representing the mean *±* one standard deviation.

For comparison, we train cryoDRGN for the same duration using the same single GPU. We use the default settings of cryoDRGN, as detailed in Appendix A.2.

We plot the mean of the FSC curves between the predicted volumes and the corresponding ground truth volumes in Figure 6, for both cryoSPHERE and cryoDRGN. This plot shows that we perform better than cryoDRGN at both the 0.5 and 0.143 cutoffs. There are two reasons behind this.

First, we fit our base structure into a volume reconstructed from a homogeneous reconstruction method. This aligns the non-moving parts of the protein with the right places, as provided by the dataset. Without this step, it is likely that some medium-scale part of the protein would have been misplaced, potentially reducing the performance of our algorithm at the 0.5 cutoff; see Appendix A.2 for evidence of this. It is important to note, that this step significantly improves our results, and it comes at almost no extra cost since it relies on the backprojection algorithm.

Secondly, deforming a good AlphaFold structure offers finer details than cryoDRGN. This accounts for the better performance of cryoSPHERE over cryoDRGN at the 0.143 cutoff. Refer to Appendix A.2 for examples of reconstructed structures and volumes.

Moreover, our method demonstrates robustness to a limited number of images. Even with only one pose per structure (totalling 10^4^ images), cryoSPHERE maintains similar performance, while cryoDRGN suffers a significant drop in the FSC curves. Further details can be found in Appendix A.2.

## 5. Discussion

CryoSPHERE presents several advantages compared to other methods for volume and structure reconstruction:

### Efficiency in Deformation

Deforming a base structure into a density map avoids the computationally expensive 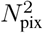 evaluation required by a decoder neural network in methods implicitly parameterising the grid, such as Zhong et al. (2021a); Levy et al. (2022). The neural network size is vastly reduced compared to state of the art methods, which explicitly parameterise the density grid of size 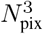, see e.g. Punjani & Fleet (2023). Furthermore, direct deformation of a structure directly avoids the need for subsequent fitting into the recovered density map.

### Reduced Dimensionality and Noise Resilience

Learning one rigid transformation per segment, where the number of segments is much smaller than the number of residues reduces the dimensionality of the problem. This results in a smaller neural network size compared to approaches acting on each residues, such as Rosenbaum et al. (2021). Rigidly moving large portions of the protein corresponds to low-frequency movements, less prone to noise pollution than the high-frequency movements associated with moving each residue independently. In addition, since our goal is to learn one rotation and one translation per segment, a latent variable of dimension 6 × *N*_*segm*_ is, in principle, a sufficiently flexible choice to model any transformation of the base structure. Choosing the latent dimension is more difficult for volume reconstruction methods such as (Zhong et al., 2021a).

### Interpretability

CryoSPHERE outputs segments along with one rotation and one translation per segment, providing valuable and interpretable information. Practitioners can easily interpret how different parts are moving based on the transformations the network outputs. This interpretability is often challenging for deep learning models such as Zhong et al. (2021a); Rosenbaum et al. (2021).

Section 4 demonstrates cryoSPHERE’s capability to recover continuous and discrete heterogeneity while performing structure reconstruction. It also demonstrate that even though the choice of *N*_*segm*_ impact the FSC to the ground truth, cryoSPHERE is able to recover the correct motion for the entire range of *N*_*segm*_ values. It is also able to keep the minimum necessary number of domains when the user sets an unnecessarily high *N*_*segm*_, as seen in Section 4.1.

While using a base structure *S*_*0*_ provides detailed information missed by cryoDRGN, especially evidenced by our better 0.143 cutoffs, it can introduce bias on medium scale elements, especially if *N*_*segm*_ is too small. We alleviate this problem by fitting S_0_ into a volume reconstructed using a simple backprojection algorithm. This, combined with cryoSPHERE, achieves better 0.5 cutoffs than cryoDRGN, showcasing its effectiveness in debiasing the results. If such a volume is unavailable, simply increasing *N*_*segm*_ can reduce the bias.

Our study opens up for significant advancements in predicting protein ensembles and dynamics, critically important for unraveling the complexity of biological systems. By predicting all-atom structures from cryo EM datasets through more realistic deformations, our work lays the foundation for extracting direct insights into thermodynamic and kinetic properties. In the future, we anticipate the ability to predict rare and high-energy intermediate states, along with their kinetics, a feat beyond the reach of conventional methods such as molecular dynamics simulations.

It would be interesting to assess how much our segmentation correlates with bottom-up segmentation into domains conducted on the “omics” scale, see e.g (Lau et al., 2023). To achieve this quantitatively, we would need many examples of moving segments from cryo EM investigations to match the millions of segments from the “omics” studies. Therefore, we leave this investigation to later work.

Finally, this work is an important milestone in showing that one can learn a segmentation of the protein that is intimately linked to the change of conformation of the underlying protein, in an end-to-end fashion. This is in contrast to other works that perform the segmentation based on residue distances of the underlying atomic model only, usually not in an end-to-end way. The application of cryoSPHERE to real dataset requires a number of careful choices that we leave to future work.

## Acknowledgement

G.D is financially supported by the Wallenberg AI, Autonomous Systems and Software Program (WASP) and the Data-Driven Life Science (DDLS) program through the WASP-DDLS collaboration.

This project would not have been possible without the computing resources provided by the Knut and Alice Wallenberg Foundation at the National Supercomputer Centre (Berzelius resource).

We also thank Claudio Mirabello at the National Bioinformatics Infrastructure Sweden at SciLifeLab for providing us access to his AlphaFold installation on Berzelius.

## Footnotes

1 https://github.com/Gabriel-Ducrocq/VAECryoEM

## Notes

### Competing Interest Statement

The authors have declared no competing interest.

### Summary of Updates

Change the order of the authors in the metadata.

