## Appendices for "cryoSPHERE: Single-particle heterogeneous reconstruction from cryo EM"

| Parameter | Value |
|---|---|
| dfU | 15301.1 Å |
| dfV | 14916.4 Å |
| dfang | 5.28 degrees |
| spherical aberration | 2.7 mm |
| accelerating voltage | 300 keV |
| amplitude contrast ratio | 0.07 |

*Table 1.* Table of the parameters used to CTF corrupt the generated images. The same values were used for all images of all datasets.

### A. Experiments

In this appendix, we provide more details on the experiments of Section 4.

In all the experiments of Section 4, each posed structure is converted into a volume, which is then projected into 2D images according to our image formation model in (2), with $\sigma = 1$. The same CTF corruption is applied with parameters specified in Table 1 and finally noise is added to achieve a SNR equal to 0.1. Here, SNR is defined as the ratio of the variance of the images to the variance of the noise.

#### A.1. Toy dataset

For this experiment, we predict the phytochrome structure using AlphaFold multimer(Evans et al., 2021) on its amino acid sequence with the UniProt (The UniProt Consortium, 2021) entry Q9RZA4. This protein forms a dimer with 755 residues on each chain. We define two domains for simulation purposes. The first domain comprises the first chain and the first 598 residues of the second chain. The second domain consists of the remaining 157 residues of the second chain. In Figure 3, we present the base structure, the decomposition into the two domains as well as the structures corresponding to the deformed base. These deformations represent the mean rotation of each mode. We rotate the second domain around the $(0, 1, 0)$ axis, sampling $10^4$ rotation angles from the Gaussian mixture:

$$0.5 \times \mathcal{N}(-\pi/3, 0.04) + 0.5 \times \mathcal{N}(-2\pi/3, 0.04) \quad (8)$$

This gives $10^4$ structures. For each image, we uniformly sample 15 rotation poses and 15 translation poses on $[-10, 10]^2$. The structure undergoes rotation, translation, and is then turned into an image according to image formation model 2. Subsequently, the images undergo CTF corruption, and noise is added to achieve $\text{SNR} \approx 0.1$. This process generates a total of 150k images, each of size $N_{\text{pix}} = 220$.

We run cryoSPHERE with $N_{\text{segm}} = 4$ for 48 hours on a single NVIDIA A100 GPU, equivalent to 779 epochs.

Due to computational constraints, the plots in Subsection 4.1 are based on only 10000 images, one per conformation.

Figure 7 illustrates the predicted angles against the latent means, demonstrating that the model effectively learns rotational motion.

Additionally, in Figure 8, we plot the distribution of predicted translation norms for the two segments, the predicted angles of rotation for segment 2 and the distribution of the dot product between the predicted axis of rotation for segment 4 and the true axis of rotation $(0, 1, 0)$. Our model predicts no translation for each segment and no rotation for the second segment. The axis of rotation is almost exactly recovered.

#### A.2. Molecular dynamics dataset

We take the structure of a phytochrome with PDB ID 4Q0J (Burgie et al., 2014) and define two domains: residues 321 to 502 of the first chain and residues 321 to 502 of the second chain. To simulate the dissociation process of the two upper domains, we perform Metadynamics(Barducci et al., 2011) simulations in GROMACS (Abraham et al., 2015; Pronk et al., 2013) with the PLUMED 2 implementation (Tribello et al., 2014). The collective variable chosen is the distance between the self-defined centers of mass (COMs) of the upper domains (residues 321-502 of chain A and B). A 100 ns simulation is conducted using the NpT ensemble, maintaining pressure control through the Parrinello-Rahman barostat. Gaussian deposition occurs every 5000 steps, featuring a height of 0.1 kJ/mol and a width of 0.05 nm. Afterwards, we extract $10^4$

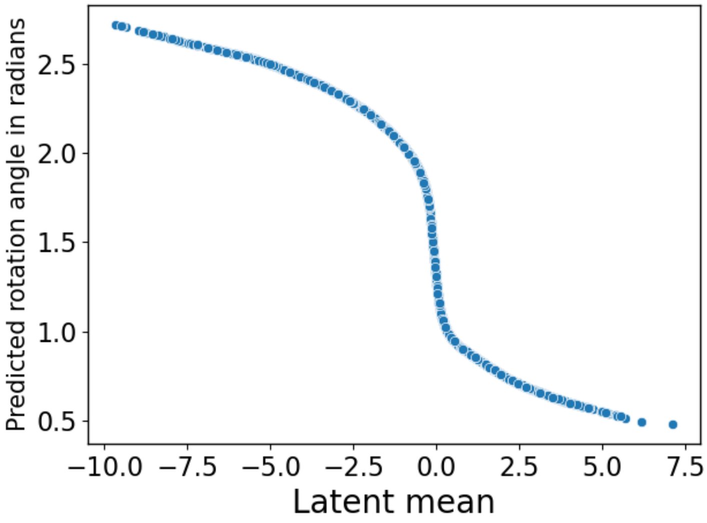


*Figure 7.* Toy dataset. Predicted angle against latent mean for cryoSPHERE. Note that for clarity $0.3$ percent of the points were removed.

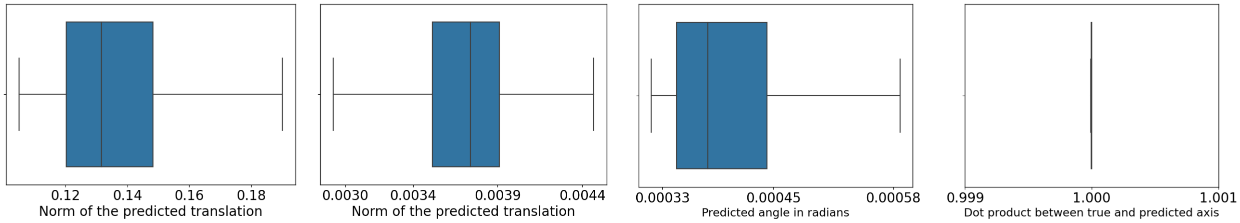


*Figure 8.* Toy dataset. Leftmost: Boxplot of the norm of the translations predicted for the fourth segment. Middle left: Boxplot of the norm of the translations predicted for the second segment. Middle right: Boxplot of the predicted angle of rotation for the second segment, in radians. Rightmost: Boxplot of the dot product between the predicted axis of rotation for the fourth segment and the true angle of rotation. CryoSPHERE recovers the right axis of rotation almost perfectly.

structures along the dry trajectory. See Figure 9 for examples of structures. From this, two datasets are generated.

The first dataset consists $15$ rotation poses uniformly sampled per structure. Translation poses are neglected in this data set, resulting in a total of $150$k images.

The same process is followed to generate the second dataset, with the exception that only one rotation pose per structure is assumed. This results in $10$k images.

For both datasets, we use $N_{\text{pix}} = 190$, and each pixel is of size $1$Å. The encoder is a 4-hidden-layer neural network with fully connected hidden layers of dimension of $2048, 1024, 512, 512$. The decoder is a 2-hidden-layer neural network with fully connected hidden layers of dimension $350, 350$. We set the batch size to $100$ and use the ADAM optimizer (Kingma & Ba, 2017) with a learning rate of $0.0003$, unless otherwise stated. The latent dimension is set to $40$ with $N_{\text{segm}} = 6$ for all runs, unless stated otherwise. For the 150k dataset, we train for 24 hours on a single NVIDIA A100 GPU, corresponding to 642 epochs. The 10k dataset is trained for 12 hours on the same single GPU, corresponding to 4114 epochs.

To compare cryoSPHERE to cryoDRGN, we train cryoDRGN using default settings - a batch size of 8 and a latent dimension equal to 8 - for the same duration, on the same single GPU used for our algorithm. To maintain consistency when comparing volumes generated from cryoDRGN, our methods, and the ground truth, we convert the structures (both ground truth and predicted with cryoSPHERE) into volumes using the same image formation model employed to generate the dataset.

For computational efficiency, the FSC plots and distances plots are not based on all structures. In the case of the 150k dataset, only one image per structure is used to compute the distances. All images in the 10k dataset are utilized. Consequently, the distance plots are based on 10k images in both cases. For the computation of the FSC curves, we select only 1000 images

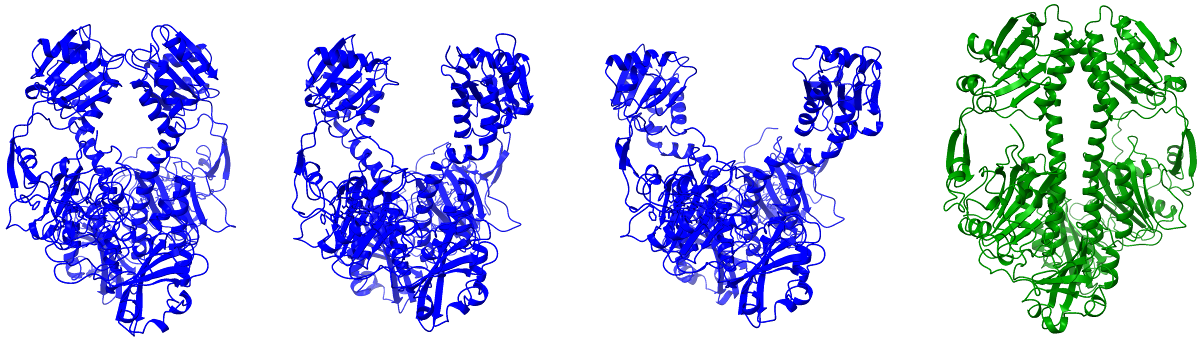

*Figure 9.* From left to right. 1/ The starting close conformation of the MD simulation. 2/ A medium-open structure arising from the MD simulation. 3/ An open conformation towards the end of the MD simulation. 4/ The AlphaFold structure used as a base structure.

evenly distributed among these 10000 structures.

##### A.2.1. 150K DATASET

Now, we test the robustness of our method to the choice of a base structure.

First, we perform two runs with $N_{\text{segm}} = 6$. For the first run, we use the same AlphaFold structure as in section 4.3, before it was fitted into the volume obtained by backprojection. In the second run, we use the same AlphaFold structure but fitted as described in Section 4.3.

Figure 10 displays the results for the unfitted run. Despite cryoSPHERE showing excellent agreement with the ground truth in terms of distances compared to cryoDRGN, it misses some medium scale elements, while maintaining a better level of small scale details. We observe underperformance at the $0.5$ cutoff but outperformance at the $0.143$ cutoff. For example, considering the ground truth volumes of the first MD structure and the corresponding reconstructed volumes from cryoSPHERE and cryoDRGN in Figure 13, we note that when only using an AlphaFold structure without postprocessing, our method can be slightly off on some medium-scale elements, especially at the bottom of the protein. As hypothesizing $N_{\text{segm}} = 6$ is not sufficient to break these elements in specific segments, cryoSPHERE does not move them, retaining the features of the AlphaFold structure, potentially misplaced. In contrast, cryoDRGN seems to be closer to the ground truth on these elements. However, upon closer inspection of the reconstructed volumes, we observe that cryoSPHERE gives much finer details throughout the volume, a charecteristic not replicated by cryoDRGN.

Figure 11 showcases the results for the fitted run. CryoSPHERE now outperforms cryoDRGN at both the $0.5$ and $0.143$ cutoffs while maintaining an excellent agreement with the ground truth. The fitting of the AlphaFold structure has a significantly positive impact on the performances of our method.

For both runs, we compare the reconstructed volumes of the same image from our method to the ground truth in Figure 14. We observe that throughout the volume, the parts that were off compared to the ground truth with the unfitted AlphaFold structure are much better placed with the fitted AlphaFold structure.

Next, we conduct two additional runs with the 150k dataset: one using the first structure from the MD simulation and the other using the last structure of the MD simulation. Figure 16 displays the results for the first MD structure as base structure, showcasing an excellent agreement with the ground truth. In addition, the FSC curves experience a drop compared to the fitted AlphaFold structure, but they are similar to the unfitted AlphaFold structure. This is likely because using a base structure belonging to the dataset introduces less bias than using an AlphaFold-generated structure with no fitting. Figure 17 presents the same plot, starting from the last MD structures. The conclusions are the same, although we tend to overestimate the distances to some extent. However, the FSC curves are significantly improved and now match the fitted AlphaFold structure run.

Figure 18 shows the segment decompositions recovered by cryoSPHERE for all these runs, revealing their overall similarity. Notably, two small segments are identified in the helices linking the upper part to the lower one. This is not problematic since our decomposition is soft, allowing for smooth transitions between no motion and the motion of the upper domains.

As expected, the quality of the results depends on the quality of the base structure used. Fortunately, AlphaFold can provide

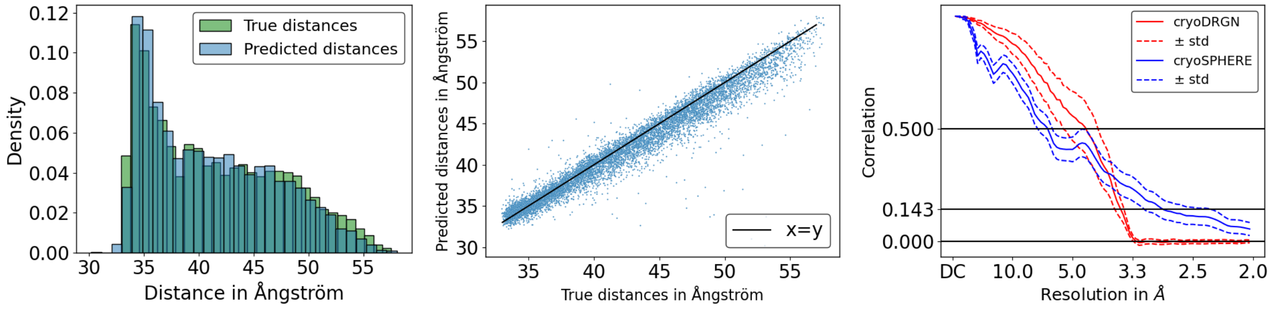


*Figure 10.* MD 150k dataset unfitted AlphaFold structure. Left: Histograms of the distances between the two upper domains. The true distances are in green. The recovered distances are in blue. Middle: Predicted against true distances in Ångström. The black line corresponds to $x = y$. Right: Plot of the Fourier shell correlation curves. The mean across structures for cryoSPHERE is in plain blue. Dotted blue represents the mean $\pm$ one standard deviation. The red, plain line represents the mean of cryoDRGN across volumes, with the dotted red line representing the mean $\pm$ one standard deviation.

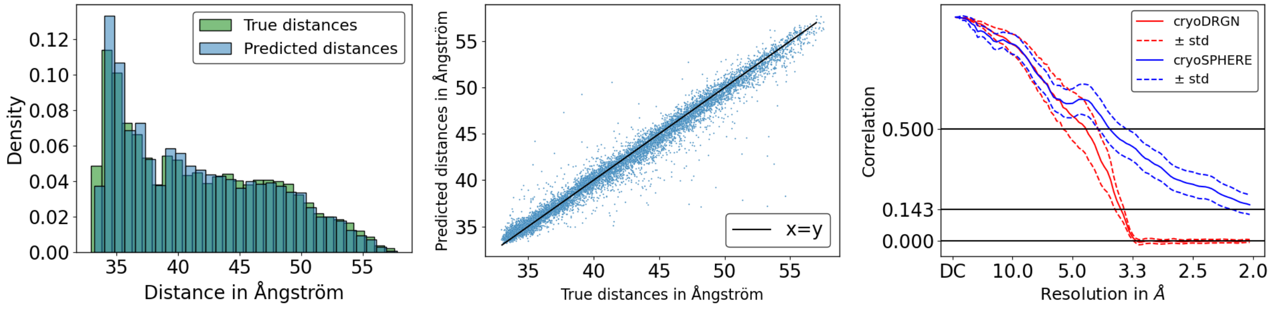


*Figure 11.* MD 150k dataset fitted AlphaFold structure. Left: Histograms of the distances between the two upper domains. The true distances are in green. The recovered distances are in blue. Middle: predicted against true distances in Ångström. The black line corresponds to $x = y$. Right: Plot of the Fourier shell correlation curves. The mean across structures is in plain blue. Dotted blue represents the mean $\pm$ one standard deviation. The red, plain line represents the mean of cryoDRGN across volumes, with the dotted red line representing the mean $\pm$ one standard deviation.

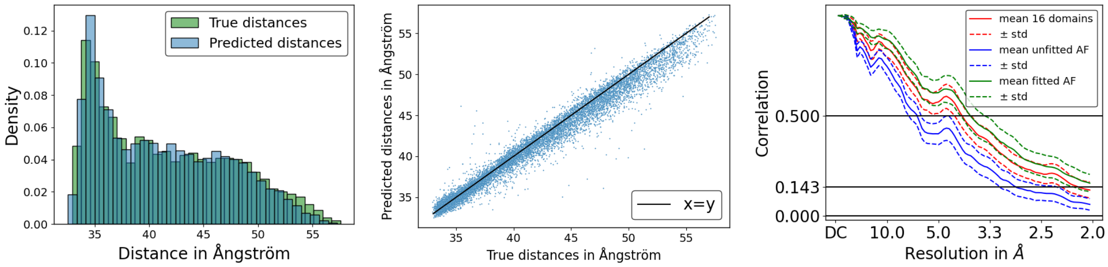


*Figure 12.* 150k dataset with the unfitted AlphaFold structure and 16 segments. Left: Histograms of the distances of the two upper domains. The true distances are in green. The recovered distances are in blue. Middle: predicted against true distances in Ångström. The black line corresponds to $x = y$. Right: Fourier shell correlation curves for cryoSPHERE. The mean of of the unfitted AlphaFold with 6 segments is in blue. Dotted blue represents the mean $\pm$ one standard deviation. The red, plain line represents the mean of the unfitted AlphaFold run with 16 segments, with the dotted red line representing the mean $\pm$ one standard deviation. The green, plain line represents the mean of the run with the fitted AlphaFold structure and 6 segments.

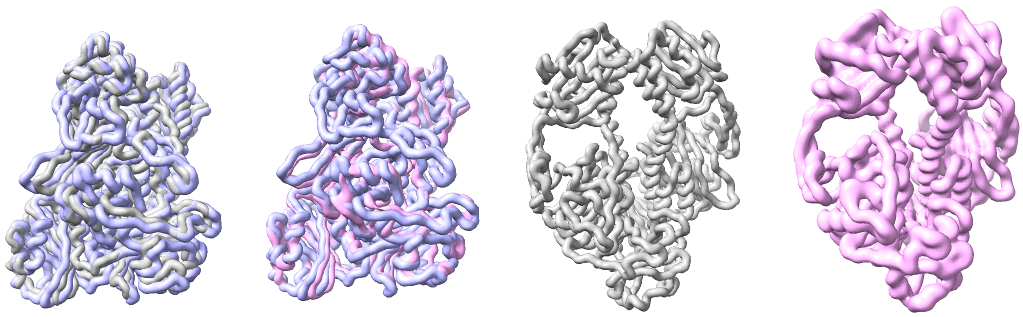

*Figure 13.* Left: comparison of a ground truth volume and the corresponding reconstructed volume from cryoSPHERE. Gray is cryoSPHERE, purple is the ground truth. Middle left: comparison of a ground truth volume and the corresponding reconstructed volume from cryoDRGN. Pink is cryoDRGN, purple is the ground truth. Middle right: cryoSPHERE. Right: cryoDRGN.

reasonable structures that we can further fit into density maps obtained with traditional methods. This approach combines the strength of both world: the more static parts of a protein can exhibit good resolution with traditional tools, and we can fit in the corresponding part of an AlphaFold structure, thereby debiasing the method to some extent. On the more mobile parts where traditional methods may have lower resolution, AlphaFold can provide improved resolution.

In a final experiment, we attempt to increase the number of segments to $N_{\text{segm}} = 16$ using the unfitted AlphaFold structure as the base structure. We run our algorithm for 24 hours on the same single GPU used in previous experiments. Figure 12 demonstrates an excellent agreement with the ground truth in terms of distances. The FSC curves indicate that increasing the number of segments actually improves the results, approaching the performance of the run with the fitted AlphaFold structure, despite using the unfitted AlphaFold structure. Figure 15 visually confirms this improvement. As expected, more segments allow the model to adjust better to the data, reducing the bias introduced by the AlphaFold structure. However, this improvement comes at a cost: in 24 hours, cryoSPHERE with the unfitted AlphaFold structure and 6 segments makes 642 epochs, whereas the run with 16 segments completes 421 epochs.

##### A.2.2. 10K DATASET

To assess the performance under data conditions, we run our method with only $10^4$ images, $N_{\text{segm}} = 6$ and the same unfitted AlphaFold base structure and settings as in subsection A.2.1. We train our network and cryoDRGN for only 12 hours only.

Figure 19 demonstrates that cryoSPHERE maintains an excellent agreement with the ground truth and is not significantly impacted by the scarcity of data. On the contrary, the FSC curves indicate that the performance of cryoDRGN is affected. This is further confirmed by Figure 20, where cryoSPHERE has roughly the same FSC curves between the 10k and 150k datasets, while cryoDRGN experiences a significant drop in performance. This is likely because the AlphaFold structure

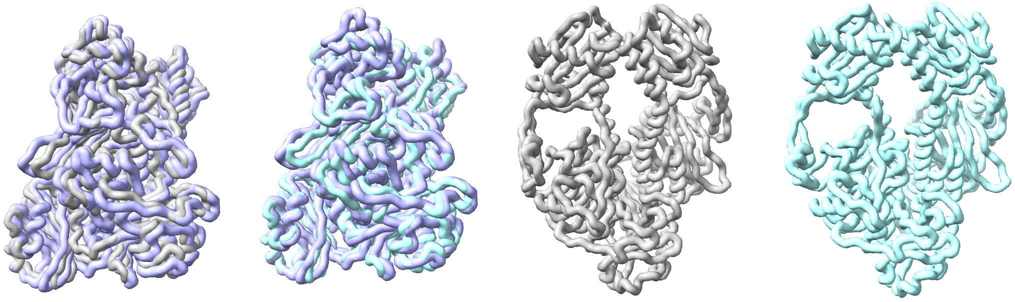

*Figure 14.* left: comparison of a ground truth volume and the corresponding reconstructed volume from cryoSPHERE with the unfitted AlphaFold structure. Gray is cryoSPHERE, purple is the ground truth. Middle left: comparison of a ground truth volume and the corresponding reconstructed volume from cryoSPHERE with the fitted Alphafols structure. Blue is fitted, purple is the ground truth. Middle right: cryoSPHERE without fitting. Right: cryoSPHERE with fitting.

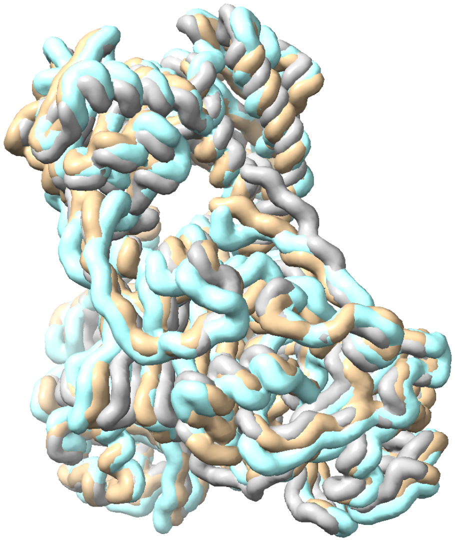

*Figure 15.* Comparison of volumes for an image of the MD 150k dataset. Gray is cryoSPHERE with unfitted AlphaFold structure and 6 segments. Tan is the same with 16 segments. Blue is ground truth. Having 16 segments leads to a volume matching the ground truth much better than 6 segments. This is true for medium scale elements throughout the volume, even at the bottom of the protein, which tends to stay still. Have a look at the coil on the middle right of the figure. Having 16 segments allows the method to break this coil into a segment and to adjust its position. Which is not the case with only 6 segments.

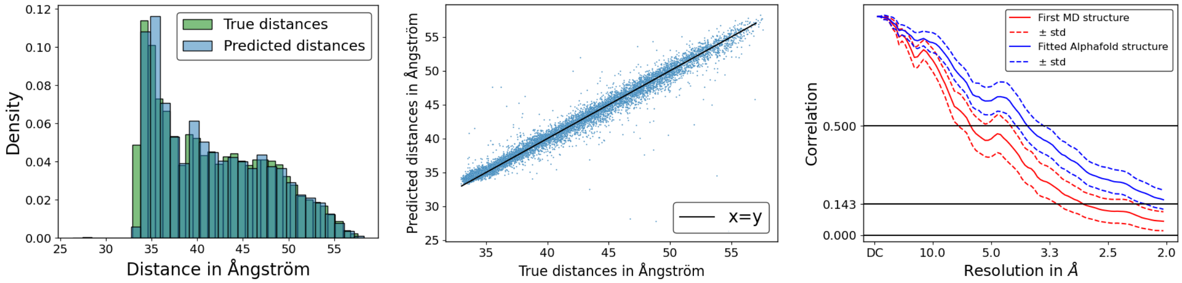


*Figure 16.* Plots for the MD 150k dataset starting from the first MD structure. Left: histograms of the true and predicted distances. Middle:predicted against true distances. Right: mean FSC curves for the AlphaFold base structure and the first MD structure as a base structure.

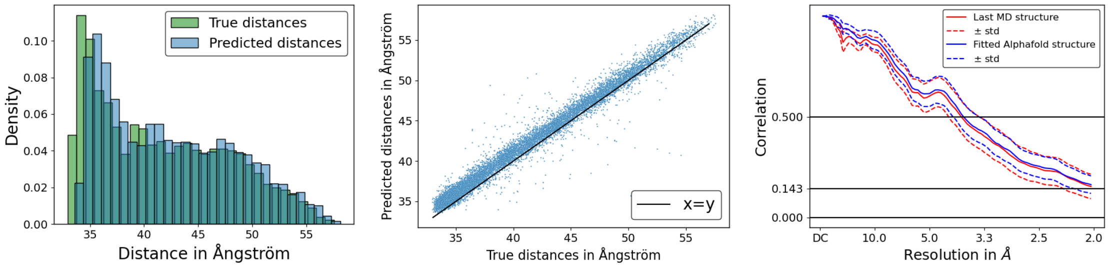


*Figure 17.* Plots for the MD 150k dataset with the last MD structure. Left: histograms of the true and predicted distances. Middle: Predicted against true distances. Right: Mean FSC curves for the AlphaFold base structure and the last MD structure as a base structure.

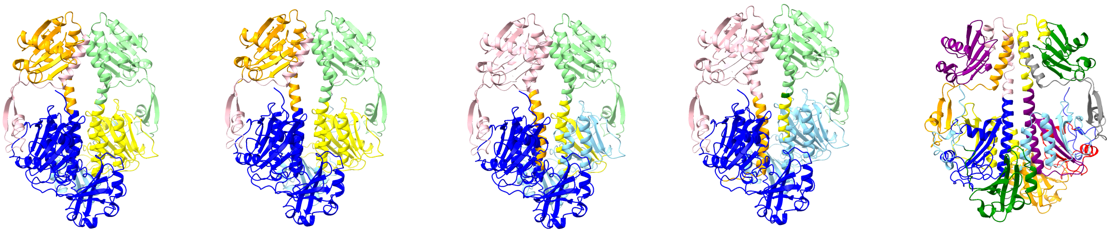

*Figure 18.* Examples of recovered segments. From left to right: 10k dataset, 6 segments. 150k dataset unfitted AlphaFold structure, 6 segments. 150k dataset with the first MD structure as the base structure, 6 segments. 150k dataset with the last MD structure as the base structure, 6 segments. 150k dataset unfitted AlphaFold structure, 16 segments.

provides strong prior knowledge.

Additionaly, we plot the recovered segments in Figure 18. Despite having fewer images, our method succesfully divides the protein into meaningful parts for the given task.

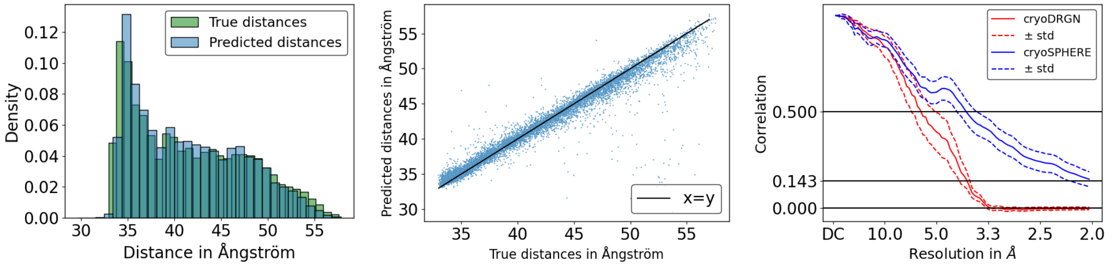


*Figure 19.* For the MD 10k dataset. Left: Histograms of the true and predicted distances. Middle: Predicted against true distances. Right: mean FSC curves for the cryoSPHERE with the same base AlphaFold structure as the 150k dataset and cryoDRGN ran on this 10k dataset. The blue lines are for cryoSPHERE, the red lines for cryoDRGN.

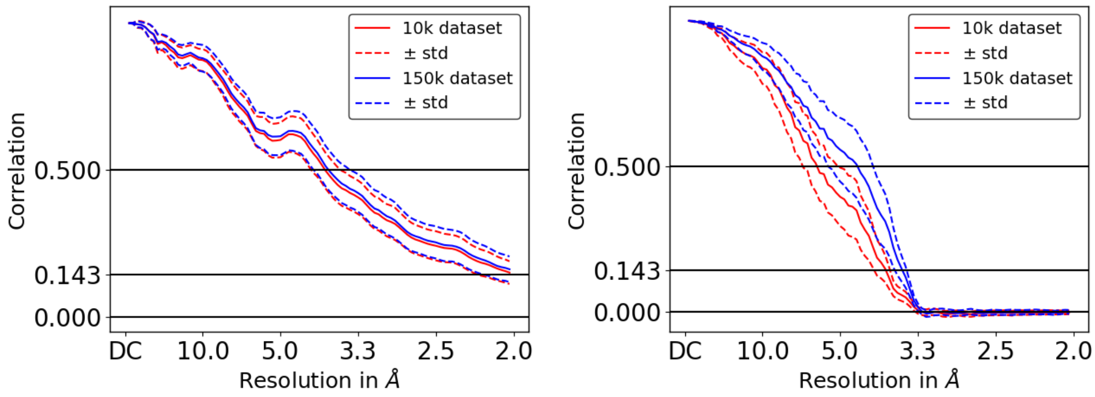


*Figure 20.* For the MD 10k dataset. Left: comparison of the FSC curves for cryoSPHERE on the 150k and 10k datasets. Right: comparison of the FSC curves for cryoDRGN on the 150k and 10k datasets. Blue represents the 150k results and red the 10k results.
